# A Ratiometric Nonfluorescent CRISPR Assay Utilizing Cas12a-Induced Plasmid Supercoil Relaxation

**DOI:** 10.1101/2023.12.04.570032

**Authors:** Noor Mohammad, Logan Talton, Selen Dalgan, Qingshan Wei

## Abstract

Most CRISPR-based biosensors rely on labeled reporter molecules and expensive equipment for signal readout. A recent alternative approach quantifies analyte concentration by sizing native λ DNA reporters, providing a simple and label-free solution for ultrasensitive detection. However, this method faces challenges in accurately quantifying size reduction of long DNA reporters via gel electrophoresis due to the sensitivity of DNA band shift to other interferences such as gel distortion. To overcome these limitations, here we developed a simple and robust ratiometric signaling strategy using CRISPR-Cas12a-induced supercoil relaxation of dsDNA plasmid reporters. In the presence of target, we observed that the fraction of supercoiled plasmid DNA decreased, and the amount of relaxed conformation (circular) increased over time. The relative percentage of supercoiled DNA to the relaxed circular DNA was analyzed by gel electrophoresis to generate an intensity-based ratiometric signal for more accurate target concentration quantification. This simple and inexpensive method is ∼100 times more sensitive when compared with the typical fluorescent reporter system. This self-referenced strategy solves the potential application limitations of previously demonstrated DNA sizing-based CRISPR-Dx without compromising the sensitivity. Finally, we demonstrated the applicability of ratiometric sensing strategy using model DNA targets such as AAV and HPV 16, highlighting its feasibility for point-of-use CRISPR-Dx applications.

**Table of Contents (TOC):** 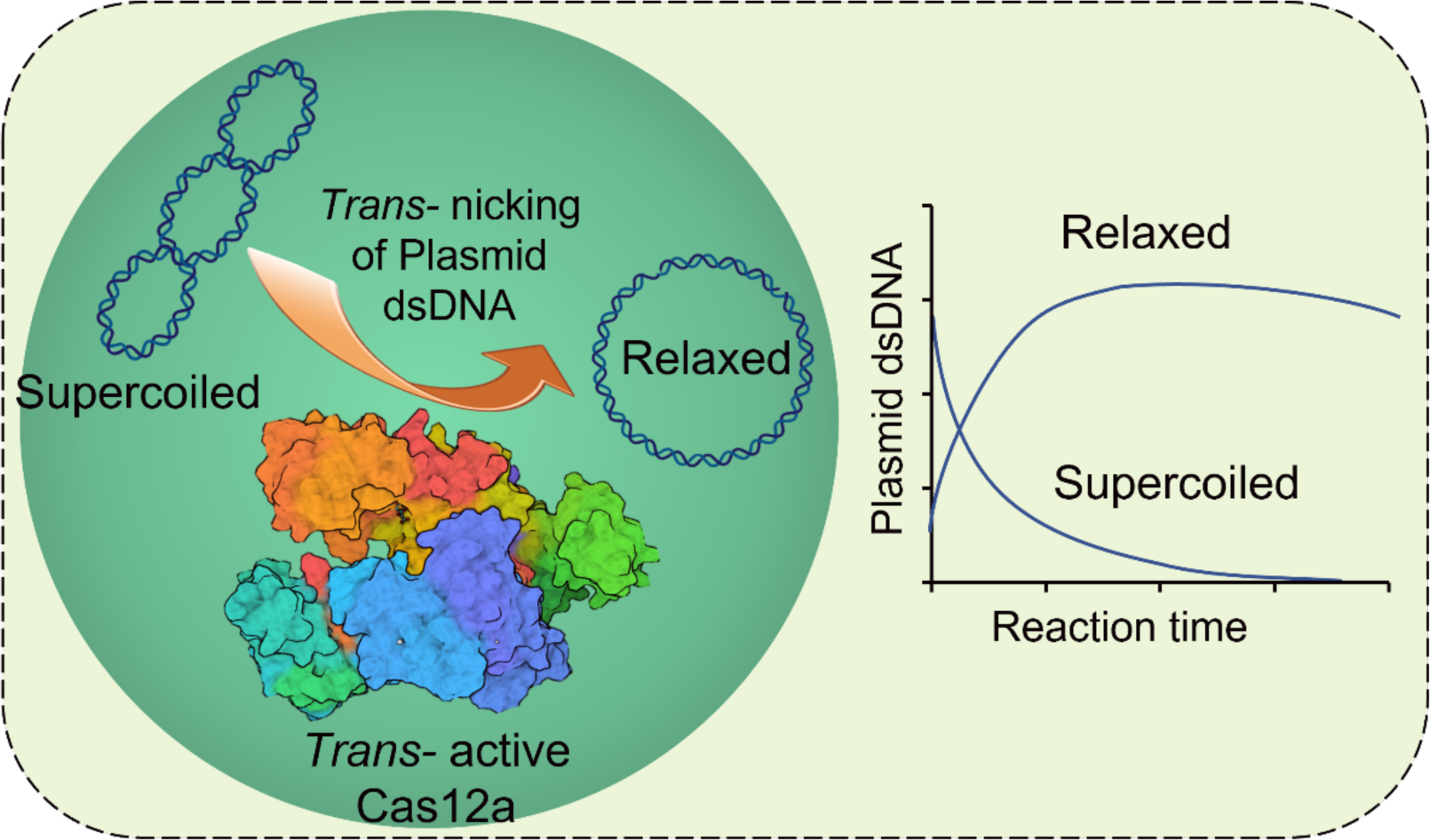

## Introduction

Rapid, inexpensive, and sensitive nucleic acid detection methods are desired for early and point-of-care (POC) diagnostics. Currently, widely applied nucleic acid detection methods, including polymerase chain reaction (PCR) assays^1–4^ and DNA sequencing^5^, are time-consuming, inaccessible for field use, and dependent on expensive instruments and skilled technical staff^6^. Besides, the current gold standard nucleic acid amplification method, PCR, faces trouble detecting small nucleic acid biomarkers such as microRNA (miRNA), as the primer design of PCR makes it infeasible to bind short nucleic acid targets (20 nts or shorter)^7^.

CRISPR (<u>C</u>lustered <u>R</u>egularly <u>I</u>nterspaced <u>S</u>hort <u>P</u>alindromic <u>R</u>epeats) with associated protein (aka CRISPR-Cas) initially appeared as a genome editing tool^8–11^ and now has evolved into a promising diagnostic tool^12–16^. For diagnostic purposes, CRISPR could be a viable alternative to PCR or DNA sequencing because CRISPR-based diagnostics (CRISPR-Dx) is isothermal, fast, sensitive, specific, and simple^17^. CRISPR-Dx came into the limelight after the discovery of Cas13a-based Specific High-Sensitivity Enzymatic Reporter UnLOCKing (SHERLOCK) platform for detecting RNA sequences^13^, and Cas12a-based DNA endonuclease-targeted CRISPR *trans* reporter (DETECTR) mechanism to detect DNA targets^12^. Numerous CRISPR-Dx platforms have been developed to date, showcasing the immense potential of CRISPR systems in addressing the longstanding challenge of achieving highly sensitive and specific nucleic acid detection through a comparatively straightforward process^14,15,18–48^. CRISPR-Cas12a particularly draws extensive attention, given its ease of reconfiguration for detecting a wide range of human, animal, and plant diseases^12,19,22,25,49–52^, including gene mutations^34,53^.

For signal acquisition purposes, CRISPR-Cas12a-Dx typically relies on fluorescent, colorimetric, or electrochemical readout^12,13,18,23,29,31,33,34,36,41,47,50,54–57^. The signal is mainly achieved when CRISPR-Cas12a nonspecifically cleaves the reporter molecules (typically ssDNA) via its unique *trans*-cleavage mode and changes the fluorescence^12,13,18,23,31,33,34,41,47,56^, color^36,50,54,55^, or current states^29,57,58^ of the reporters. One of the major limitations of these signal generation strategies is the need for expensive labels and readout instruments. Instead, we previously demonstrated a simple, inexpensive, and nonoptical CRISPR-Cas12a-based sensing platform to detect short ssDNA at sub-picomolar detection sensitivity using ds λ DNA as a novel reporter. This sizing-based method measures the length reduction of λ DNA reporters via gel electrophoresis to quantify target concentrations without any fluorescent labels or complicated optical instrument^17^. However, precise determination of band shift for long DNA reporters (e.g., λ DNA) is challenging because not only it requires a control λ DNA reporter running on the same gel to serve as the reference, but also the band positions are subject to various interferences, such as gel uniformity and uneven electric field. In addition, the accurate band shift for long DNA (e.g., λ DNA) is very difficult to evaluate because DNA fragments >20 kb are hard to separate and run nearly at the same speed in the conventional gel electrophoresis technique^59^. Specialized gel electrophoresis (e.g., pulse field gel electrophoresis) is more efficient to separate larger DNA fragments, but is more expensive and complex^59,60^. Recently, the sizing-based detection concept has also been applied to a new class of hybrid reporters (i.e., dsDNA backbone plus a 3’ toe)^61^. Although the separation of short hybrid reporters is much faster and easier, the band distortion challenge still remains.

Here, we demonstrated a self-calibrated and nonfluorescent CRISPR-Cas12a-based sensing platform to detect short ssDNA or dsDNA utilizing the conformation change of supercoiled plasmid DNA reporters to generate ratiometric signal output without the need for additional molecular reference. Different from the previous sizing approach that relies on precise band position determination, this new methodology produces ratiometric signals based on band intensities, eliminating many gel distortion-induced interferences. Three circular dsDNA plasmids, namely pUC19 (2.69 kb)^62^, pBR322 (4.36 kb)^63^, and ΦX174 (5.37 kb)^64^ were employed as novel ratiometric reporters for the CRISPR-Cas12a assay. After the CRISPR reaction, we observed the conformation change of the dsDNA plasmid reporters from supercoil to relaxed circular form induced by *trans*-active LbCas12a. We suspect that target-activated Cas12a nicks and relaxes supercoiled plasmids via its *trans*-cleavage mode, producing first circular and then linear forms of plasmids. The extent of conformation change (supercoil to circular) induced by *trans*-active Cas12a is proportional to the target concentration. By analyzing the relative population of supercoiled and circular plasmid reporters, we achieved ultrasensitive detection of ssDNA targets with a limit of detection (LOD) of ∼250 fM, superior to fluorescent reporter-based systems by ∼100 times. Furthermore, we demonstrated the application of this self-referenced approach for detecting real-life DNA targets such as adeno-associated virus (AAV) and human papillomavirus 16 (HPV16).

## Results and Discussion

### Discovery of the dsDNA plasmid reporter

CRISPR-Cas12a-based diagnostic schemes have mainly been demonstrated based on the *trans*- cleavage of ssDNA reporter by Cas12a protein^12,15,24,31,36,41^. Recently, we discovered that CRISPR- Cas12a can also actively *trans*-cleave ds λ DNA substrate^17^. Although it is still unclear how ds λ DNA gets *trans*-cleaved, the use of dsDNA reporters would not only enrich the varieties of CRISPR-Dx, but it could also overcome the limitations of the typical fluorescence-based reporting system, such as the need for fluorescent labeling and readout. To further explore the *trans*-cleavage behavior of Cas12a on other dsDNA substrates, we chose to screen a set of plasmid DNA molecules. Plasmids are small, circular, dsDNA molecules that can be tightly supercoiled in order to fit inside the cell. In the initial test, two model reporters: 1-kilobase pair (kb) linear dsDNA and 2.69-kb pUC19 dsDNA plasmid, were chosen and compared. To run CRISPR-Cas12a assay, a 20- nt long synthetic ssDNA was used as the artificial target (see sequence in **Table S1**). In **Figure 1a**, lanes 1 and 4 are negative controls, where 1-kb dsDNA reporter and pUC19 DNA were used, respectively, but no target was added. As expected, no reporter cleavage was observed since the target was not there to activate LbCas12a. It is worth mentioning that intact pUC19 plasmid has two distinct bands in lane 4, indicating the presence of supercoiled and relaxed (circular) conformations of pUC19. Lanes 2 and 3 were testing lanes for the 1-kb dsDNA reporter and 2.5 nM ssDNA target was present. Similarly, lanes 5 and 6 were testing lanes for pUC19 DNA reporter in the presence of targets. No collateral cleavage was observed for the 1 kb dsDNA reporter when targets were added (**Figure 1a**, lanes 2 and 3), which confirms the conventional knowledge of the field that CRISPR-Cas12a does not typically *trans*-cleave dsDNA substrates (λ DNA is an exception)^12^. However, for the pUC19 reporter, we observed a significant change in band pattern in the presence of target (**Figure 1a**, lanes 5 and 6). The supercoiled band (lower band) disappeared, the circular band (upper band) was enhanced in intensity, and finally, a new band appeared in between. We attribute this interesting pattern change to the conformation alternation of the pUC19 reporter, which underwent *supercoiled-circular-linear* transformation due to the enzymatic activity of Cas12a (**Figure 1b**). We believe the following steps occur in a sequence: first, *trans*-active Cas12a would preferably nick the twisting knots of supercoiled pUC19 (we call it *trans-*nicking) and relax pUC19 into the lower energy circular form (or relaxed form). Next, excess target-activated Cas12a could further *trans*-degrade circular pUC19 into linear fragments by completely breaking the previous nicking site (**Figure 1b**). The nicking effect of activated Cas12a was recognized in a few previous reports^65^, but it is the first time that such nicking behavior can be utilized for plasmid relaxation.

**Figure 1.**
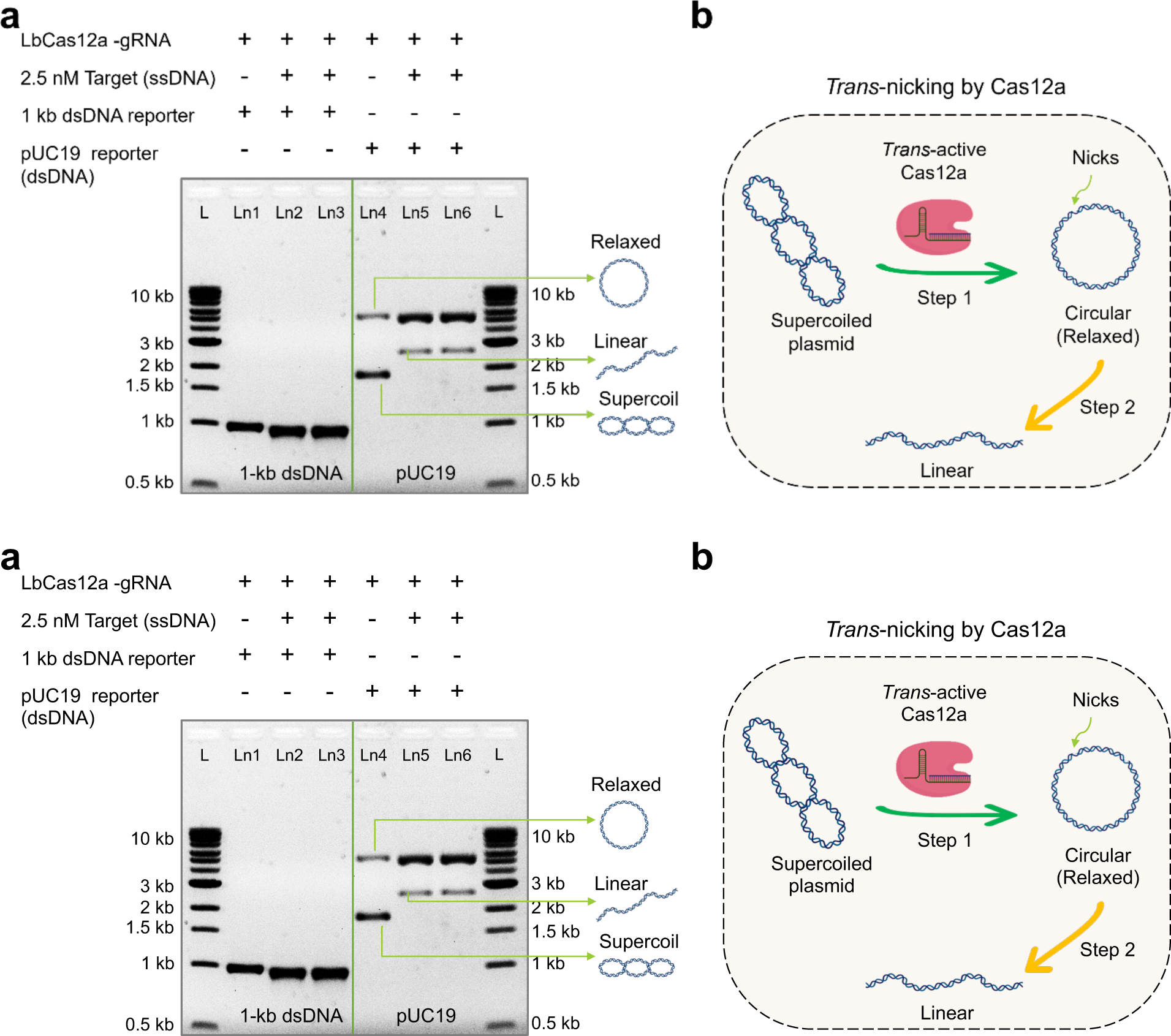
dsDNA plasmid as the CRISPR-Cas12a reporter. (a) Gel electrophoresis (1% agarose gel and 1×TBE buffer) results demonstrating the different CRISPR-Cas12a assay results of using 1-kb linear dsDNA and 2.7 kb supercoiled pUC19 dsDNA plasmid as nonspecific substrates. (b) Cartoon illustration showing the conformation change of pUC19 induced by trans-active Cas12a. The supercoiled-circular transition step is utilized for developing a ratiometric CRISPR (rCRISPR) assay later. Abbreviations: gRNA, guide RNA; nM, nanomolar; ssDNA, single-stranded DNA; kb, kilobase pair; L, 1-kb ladder; Ln, lane; TBE, Tris-borate EDTA; dsDNA, double-stranded DNA.

### Nonfluorescent ratiometric CRISPR-Cas12a assay (rCRISPR) utilizing plasmid supercoil relaxation

Next, we evaluated the feasibility of using the gel pattern change of the plasmid reporter as a new self-calibrated sensing mechanism. The developed ratiometric CRISPR assay is termed rCRISPR for simplicity. To demonstrate the nonoptical rCRISPR assay based on dsDNA supercoil relaxation, we ran CRISPR-Cas12a reaction following the standard protocol, except that we used pUC19 dsDNA as the reporter molecule. **Figure 2a** shows the schematic of rCRISPR assay using pUC19 DNA as the reporter molecule. **Figure 2b** shows the cartoon schematic illustrating the Cas12a nicking and conversion of supercoiled pUC19 into relaxed circular conformation. **Figures 2c** and **2d** denote the graphical predictions showing how the fraction of supercoiled and relaxed circular DNA would change as the CRISPR reaction proceeds at a fixed target concentration, and how the ratio of supercoiled DNA to relaxed circular DNA could change with the target concentrations at a fixed reaction time, respectively.

**Figure 2.**
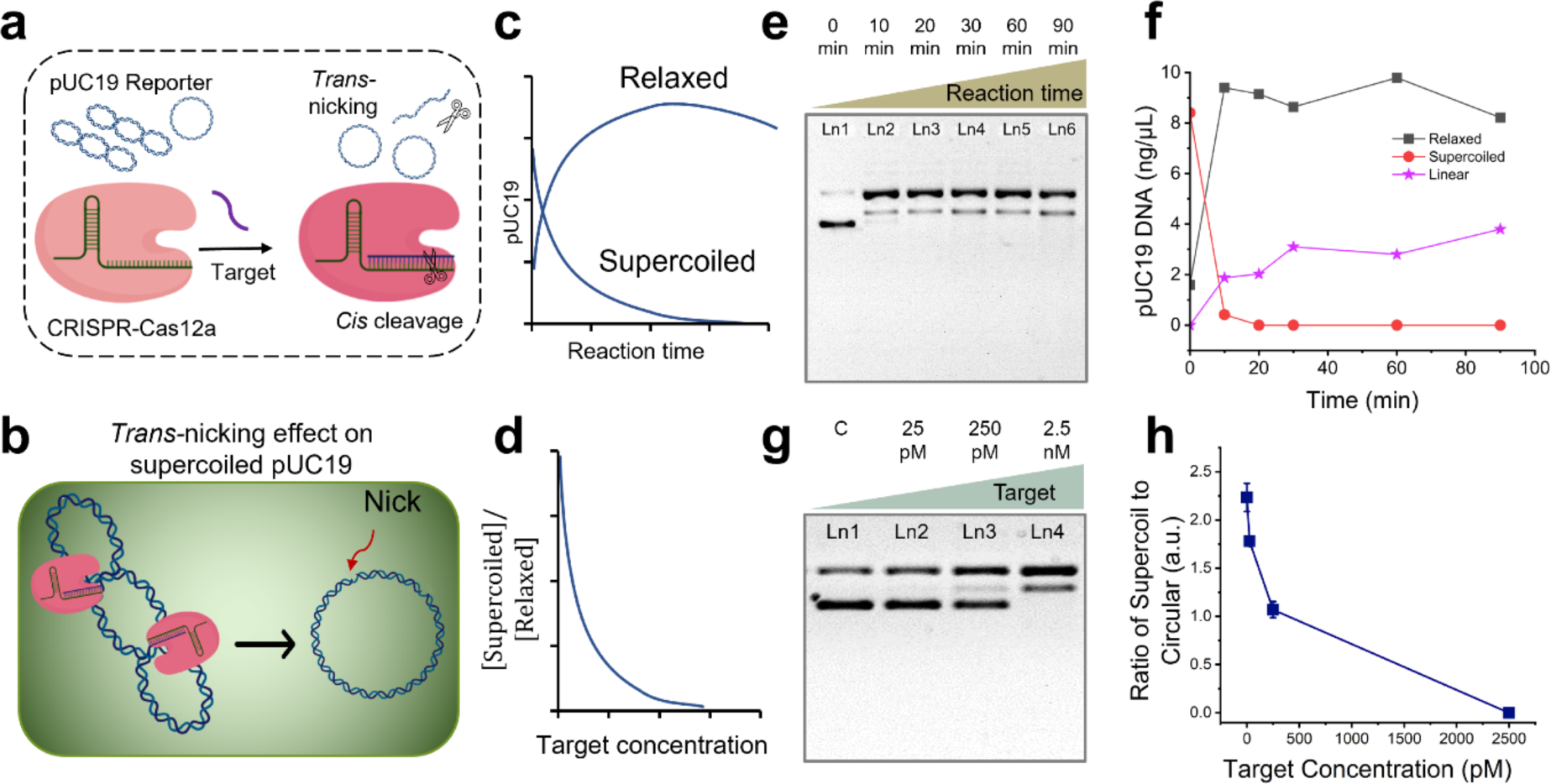
Ratiometric CRISPR-Cas12a assay using plasmid reporters. (a) Schematic of the CRISPR-Cas12a assay using pUC19 as a nonspecific substrate. (b) Close-up cartoon view showing activated Cas12a nicks the supercoil pUC19 and converts it into relaxed conformation. (c) Hypothetical graphical illustration showing how the percentage of supercoil and relaxed DNA would change as the CRISPR reaction proceeds. (d) Hypothetical graphical illustration showing how the ratio of supercoil DNA to relaxed circular DNA could change with the target concentrations. Gel electrophoresis (1% agarose gel and 1×TBE buffer) results (e) and scatter plot (f) demonstrating the nonspecific nicking of pUC19 by Cas12a over the reaction time course. DNA concentration (ng/μL) was estimated by comparing the band intensities and converted from the initial plasmid DNA concentration. Gel electrophoresis (1% agarose gel and 1×TBE buffer) results (g) and scatter plot (h) demonstrating the effect of target concentration on *trans*-nicking of pUC19 by Cas12a for a 30-min reaction. Abbreviations: c, negative control; pM, picomolar; nM, nanomolar; TBE, Tris-borate EDTA; a.u., arbitrary unit.

To verify the prediction, we ran rCRISPR reactions with pUC19 DNA reporters for 10 min, 20 min, 30 min, 60 min, and 90 min. We used 2.5 nM ssDNA target to *trans*-activate Cas12a. We observed that within 10 minutes of reaction, all supercoiled DNA vanished (aka. converted to relaxed form) (**Figure 2e**). Converting relaxed circular DNA into linear conformation might happen instantly with the supercoil relaxation; however, the linearization process was observed up to 90 minutes of reaction, as indicated by the slow darkening of the linear form band. Notably, no degradation of linear dsDNA was observed for the given reaction conditions (90 minutes of reaction, 2.5 nM target) (**Figure 2e**). The sharp slope in **Figure 2f** for supercoiled DNA confirmed that the DNA supercoil relaxation is very fast compared to the 2^nd^ step (DNA linearization) and potentially 3^rd^ step (degradation of liner DNA, which happens only for high target concentrations). This also could be the reason why we did not see much difference in gel patterns when we tested different assay times for 1 hr, 2 hrs, and 3 hrs (**Figure S1**). We believe that the *trans*-nicking of supercoil plasmids occurs even faster than *trans*-cleavage of fluorescent reporters. In the case of conventional ss fluorescent reporters, the CRISPR reaction will usually take >15 min to ensure all F-Q reporters are properly cleaved^17^ and generate detectable signals. In contrast, with similar analyte concentration (i.e., 2.5 nM ssDNA), the supercoil relaxation can be completed <10 min (**Figure 2e**) and therefore could be advantageous for developing a faster-response CRISPR biosensor.

We then performed the assay with 25 pM, 2.5 nM, 25 nM, and 0 targets (negative control) and evaluated the feasibility of ratiometric sensing for 30 min reaction time (**Figure 2g**). We divided the intensity of the supercoiled band (lower band in each lane in **Figure 2g**) by the intensity of the relaxed circular band (upper band in each lane in **Figure 2g**) to get the ratiometric intensity. We plotted the ratiometric intensity against different target concentrations, and observed that the ratiometric signal decreased as a function of target concentration (**Figure 2h**), following a similar trend to the prediction (**Figure 2d**). This initial data demonstrates the feasibility of using plasmid DNA as a novel ratiometric reporter for sensitive nucleic acid detection, which overcomes the limitations of the previously demonstrated DNA-sizing-based sensing^17^ and also excludes the need for expensive dye reporter molecules or sophisticated optical equipment.

### The universality of supercoil relaxation among different dsDNA plasmid reporters

While the *trans*-nicking of pUC19 DNA is interesting, we performed additional CRISPR reactions using a few other ds plasmids, such as pBR322 and ΦX174, to see if this supercoil relaxation mechanism is widely applicable among different plasmid DNA. **Figure 3** shows that the supercoil band (lower band) of pBR322 completely disappeared in the presence of 2.5 nM and 25 nM target concentrations (left panel, lanes 2 and 3). The same phenomenon was observed for ΦX174 (right panel, lanes 5 and 6). These results suggest that the ratiometric sensing strategy could be employed using various ds plasmid DNAs, which makes this method universal. However, the detection sensitivity would depend on the specific plasmid reporter, which is discussed in the later section.

**Figure 3.**
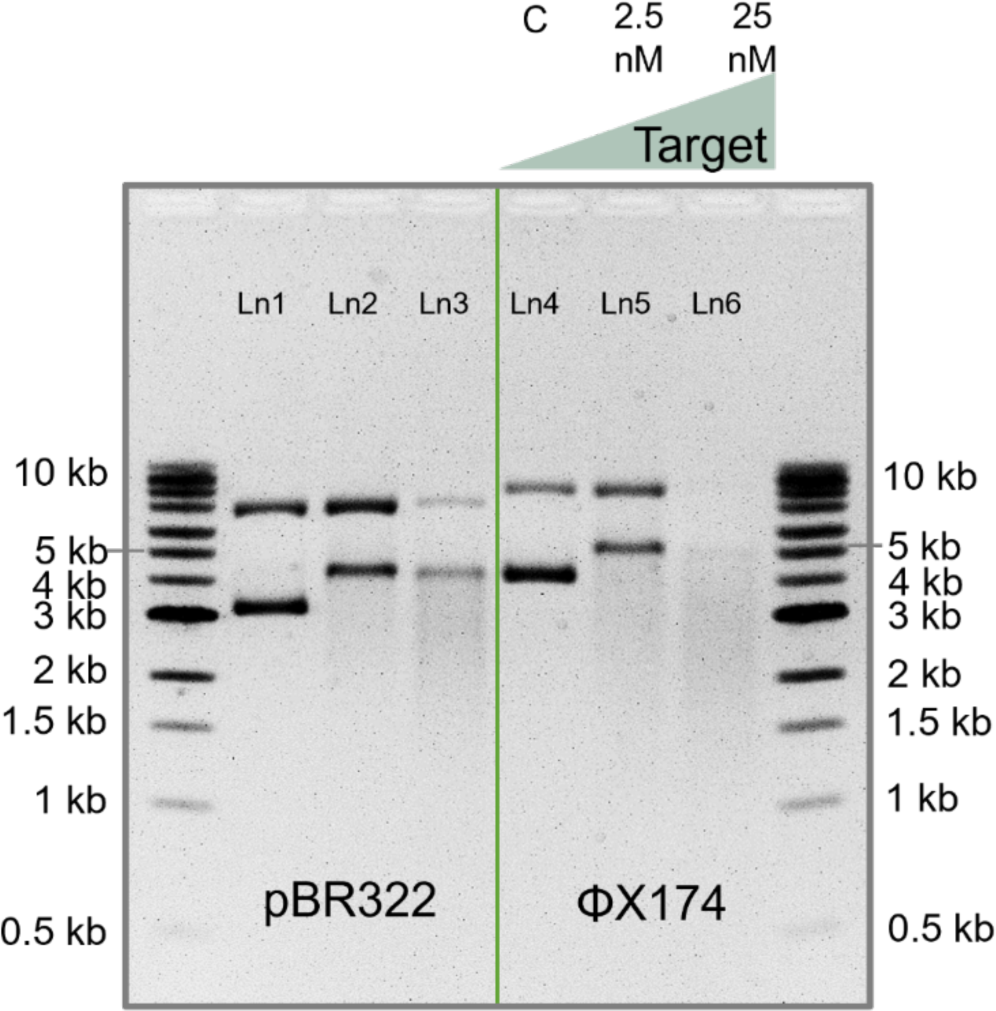
Gel electrophoresis (1% agarose gel and 1×TBE buffer) results demonstrating CRISPR- Cas12a assay using two new plasmid DNA (pBR322 and ΦX174) as nonspecific substrates. Abbreviations: nM, nanomolar; kb, kilobase pair; Ln, lane; TBE, Tris-borate EDTA.

### Assay optimization, characterization, and comparison with other reporting strategies

Among the three dsDNA plasmid reporters (pUC19, pBR322, and ΦX174), we chose pUC19 to optimize the assay conditions, and then calculated the LOD of the nonfluorescent rCRISPR assay using all three plasmid DNAs. Finally, we compared the performance of rCRISPR with the conventional fluorescent CRISPR assay based on a F-Q reporter.

We started our optimization procedure with the pretreatment of the reporter molecules. We ran rCRISPR reactions with untreated pUC19, acidic pUC19 (pH ∼6), basic pUC19 (pH ∼10), and snap-cooled^17^ pUC19. The gel images and bar charts in **Figures 4a-d** reveal that there was no effect of snap cooling on the DNA supercoil relaxation efficiency; however, acidic condition destabilized the plasmid reporter (**Figure 4b**), and basic pretreatment promoted *trans*-nicking and supercoil relaxation (**Figure 4c**). Plasmid DNA is unstable under acidic conditions because low pH promotes the nucleophilic attack of the phosphate group by water molecules, resulting in the cleavage of the phosphodiester bonds^66^, causing strand breakage and thus degrading the DNA molecule. For basic treatment, we observed that the supercoil band completely disappeared with 250 pM target concentration (**Figures 4c**, lane 3). At higher pH, the phosphate groups on the DNA backbone become deprotonated, increasing the DNA molecule’s negative charge density. This increased negative charge can repel the negatively charged phosphate groups, destabilizing the DNA helix and unwinding the ds structure^67^. This unwound ds portion of supercoil DNA could create easy access for the *trans*-active Cas12a protein.

**Figure 4.**
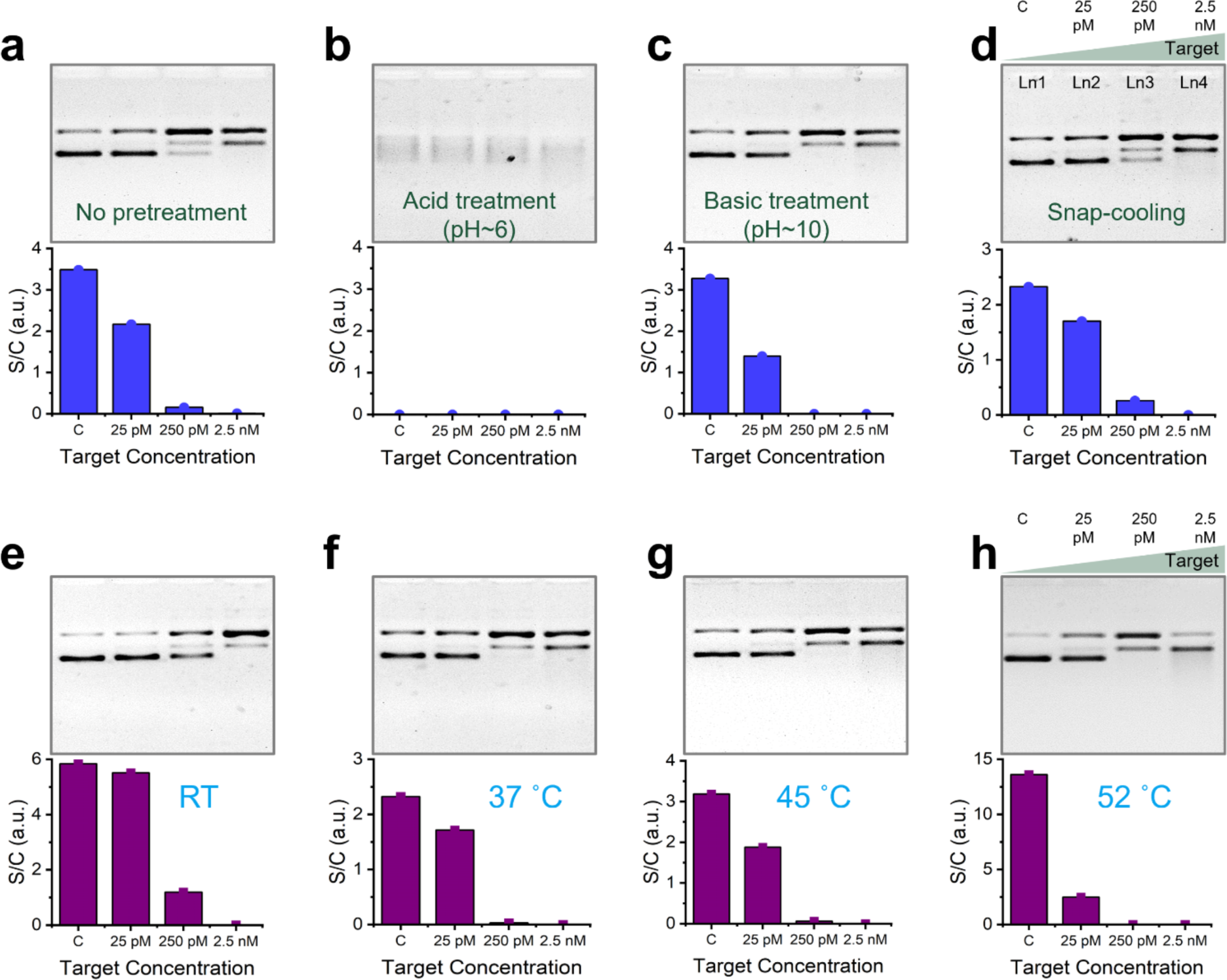
Effect of reporter pretreatment (a-d) and reaction temperature (e-h). Gel electrophoresis (1% agarose gel and 1×TBE buffer) results (top) and bar charts (bottom) demonstrating the effect of no pretreatment (a), acid pretreatment (b), basic pretreatment (c), and snap-cooling (d) on trans- nicking of pUC19. Gel electrophoresis (1% agarose gel and 1×TBE buffer) results (top) and bar charts (bottom) demonstrating the effect of reaction temperature on *trans*-nicking of pUC19 (e-h). Intensity plots of each gel have been shown in **Figure S2**. Abbreviations: c, negative control; pM, picomolar; nM, nanomolar; RT, room temperature; TBE, Tris-borate EDTA; a.u., arbitrary unit; S/C, supercoil/relaxed circular.

Figures 3e-h show the effect of reaction temperature on DNA supercoil relaxation efficiency. We observed that 250 pM target resulted in the complete disappearance of supercoil bands for all reaction temperatures except for RT (Figures 3e-h, top panels, lane 3). We concluded that RT is the least active reaction temperature among the temperatures we tested. In addition, we observed that higher reaction temperature generally favors the DNA supercoil relaxation (see results of 25 pM target in Figure 3e-h, bottom panels). However, we cannot increase the reaction temperature beyond Cas12a’s stability temperature, which is about low 50 °C. After that, Cas12a will be denatured and lose its biological function^68^.

Next, a series of reaction mixture conditions were optimized, including the concentration of ribonucleoprotein complex (RNP) and buffer composition. Figures 5a-c show the gel results (top) and quantitative bar charts (bottom) of three sets of rCRISPR reactions done with different Cas12a concentrations (e.g., 20 nM, 40 nM, and 60 nM). For each Cas12a concentration, we kept the same molar ratio of Cas12a:gRNA (2:1) according to the literature^47^. For each set, 25 pM, 250 pM, and 2.5 nM target concentrations, including a negative control, were tested. We observed that the supercoiled band (Figures 5a-c, top panels, lane 3) for 250-pM target became thinner with the increase of Cas12a concentration, which indicates that higher Cas12a concentration performed better for DNA supercoil relaxation. However, if we compare the intensity differences between negative control and 25-pM target of each set (Figures 5a-c, bottom panels), we found the maximum difference is from 20 nM Cas12a concentration. The results suggest that the optimal Cas12a concentration depends on the concentration of the target nucleic acid. In general, lower RNP concentrations are more suitable for detecting lower concentrations of target nucleic acids, while higher Cas12a concentrations are preferable for detecting higher target concentrations, as we previously observed^17^.

**Figure 5.**
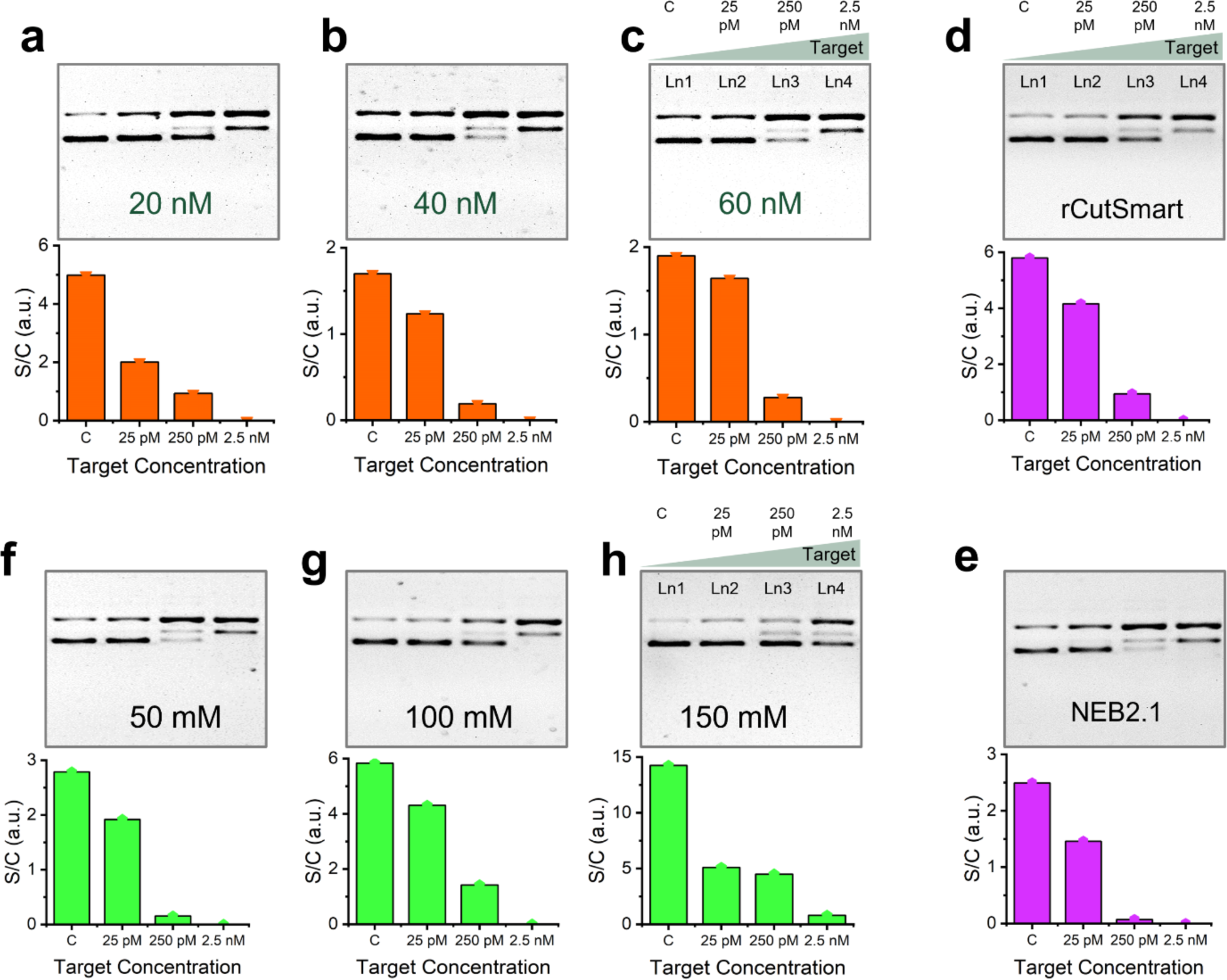
Effect of Cas12a concentration (a-c), buffer type (d-e), and salt concentration (f-h). Gel electrophoresis (1% agarose gel and 1×TBE buffer) results (top) and bar charts (bottom) demonstrating the effect of 20 nM Cas12a (a), 40 nM Cas12a (b), and 60 nM Cas12a (c) concentration; effect of rCutSmart reaction buffer (d), and NEB2.1 reaction buffer (e); effect of 50 mM NaCl (f), 100 mM NaCl (g), and 150 mM NaCl (h) on *trans*-nicking efficiency of Cas12a towards pUC19. Intensity plots of each gel have been shown in **Figure S3**. rCutSmart buffer recipe: 50 mM Potassium Acetate, 20 mM Tris-acetate, 10 mM Magnesium Acetate, 100 µg/ml Recombinant Albumin, pH 7.9@25°C. NEBuffer 2.1 buffer recipe: 50 mM NaCl, 10 mM Tris-HCl, 10 mM MgCl2, 1 mM DTT, 100 µg/ml BSA, pH 7.9@25°C. Abbreviations: c, negative control; pM, picomolar; nM, nanomolar; mM, millimolar; TBE, Tris-borate EDTA; a.u., arbitrary unit; BSA, bovine serum albumin.

Besides, we explored the effect of reaction buffer types and salt concentration (ionic strength) on DNA supercoil relaxation. We used rCutSmart buffer and NEBuffer2.1, containing two different types of salts: acetate salt vs. chloride salt (buffer compositions are mentioned in the caption of Figure 5). Both qualitative (Figures 5d-e, top panels, lane 3) and quantitative (Figures 5d-e, bar charts for 250 pM target) results suggest that NEBuffer2.1 performs better. In addition, we explored the effect of salt concentration on the *trans*-nicking of supercoil DNA (Figures 5f**- h**). We observed that higher salt concentrations (e.g., 150 mM NaCl) resulted in a lower nicking rate (Figures 5f-h, top panels, lanes 3 and 4). A higher concentration of cations such as Na^+^ is known to stabilize nucleic acid helix at hybridized state^69^. Thus, plasmids became more immune from being nicked at higher ionic strength.

After the assay optimization, we characterized the LOD of the rCRISPR assay at optimized conditions (e.g., basic pretreatment, T=52 °C, NEB2.1 buffer, etc.). We first characterized the LOD for pUC19 reporter. We titrated the assay with a broad range of analyte concentrations, ranging from 2.5 pM to 25 nM for 1 hr of reaction. The gel image is shown in Figure 6a. We observed that the supercoil band of pUC19 reporter gradually became thinner, and the relaxed circular band became stronger. Besides, a new band for linear reporter molecule started to appear with the increasing target concentrations (Figure 6a, lanes 5 and 6). We also observed that all bands vanish at 25 nM target concentration, indicating the complete degradation of the reporter molecule eventually. We plotted intensity diagrams for each lane of the gel (Figure 6b). The left peak denotes the relaxed circular form, the middle one is linear DNA, and the right peak is for supercoiled pUC19 DNA. We measured the peak intensity and plotted it in a bar chart against various target concentrations (Figure 6c). We performed statistical analysis (t-test) to determine the minimum concentration we could detect with our newly proposed ratiometric sensing strategy. We found a *p-*value less than 0.05 for all concentrations we tested in our experiment. Thus, we estimated a LOD of ∼2.5 pM for the synthetic ssDNA target based on statistical analysis.

**Figure 6.**
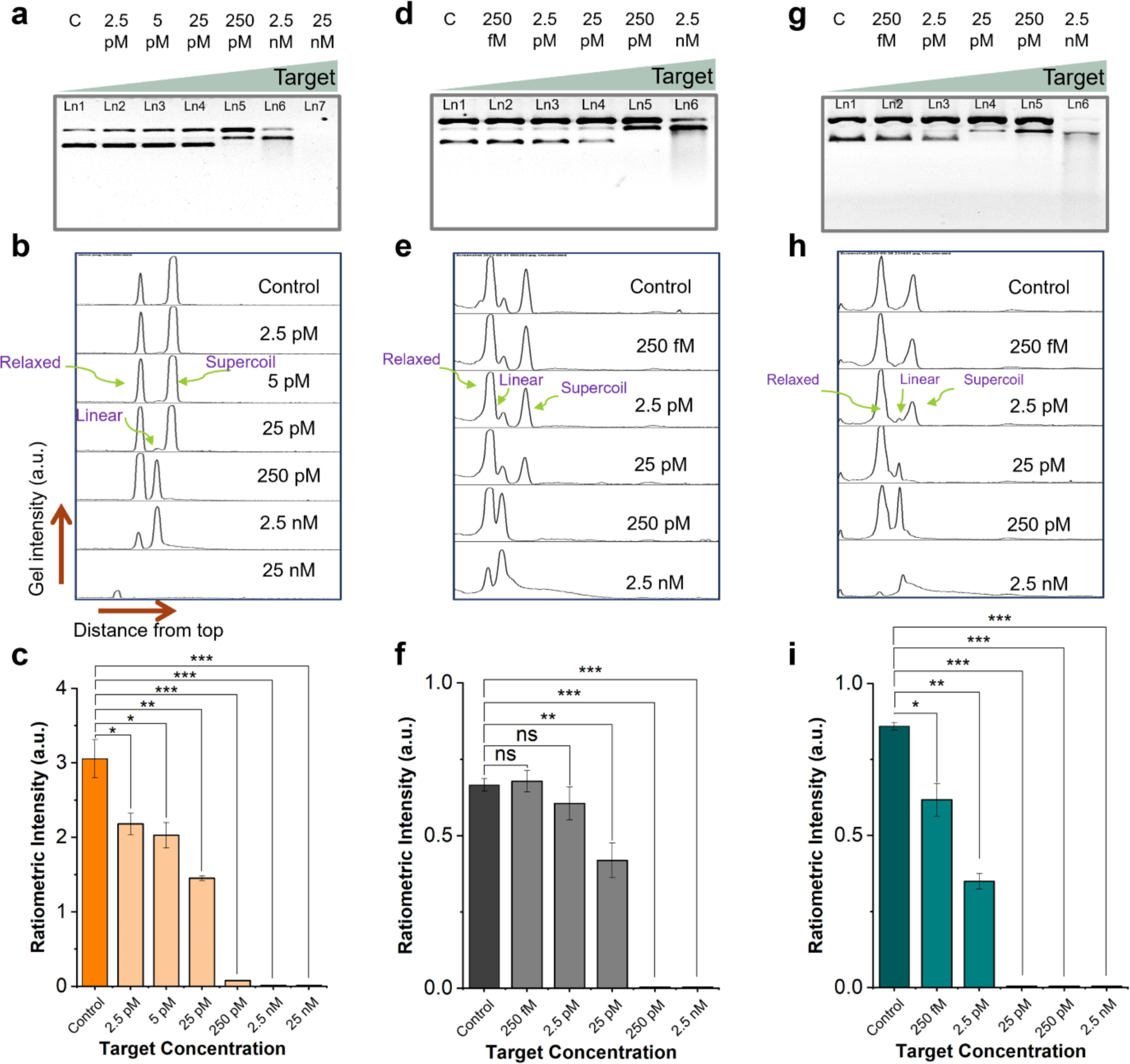
Limit of detection (LOD) determination. Gel electrophoresis (1% agarose gel and 1×TBE buffer) demonstrating the CRISPR-Cas12a induced DNA relaxation for various target concentrations using pUC19 (a), pBR322 (d), and ΦX174 (g) reporters, respectively. Gel intensity diagrams of gel lanes for pUC19 (b), pBR322 (e), and ΦX174 (h) reporters. Ratiometric intensity plotted in bar chart against different target concentrations for pUC19 (c), pBR322 (f), and ΦX174 (i) reporters. The graph shows statistical insignificance at p>0.05 (ns), significance at p<0.05 (*), p<0.01 (**), and p<0.001(***). Abbreviations: c, negative control; fM, femtomolar; pM, picomolar; nM, nanomolar; Ln, lane; TBE, Tris-borate EDTA; a.u., arbitrary unit.

Furthermore, we repeated the experiments using pBR322 (Figures 6d-f) and ΦX174 (Figures 6g-i) as the reporter molecules for varied target concentrations from 250 fM to 2.5 nM. We found LODs of 25 pM and 250 fM for pBR322 and ΦX174 reporter, respectively, using a similar analysis as mentioned above. We observed that ΦX174 outperformed all three plasmid reporters we tested. In this ratiometric sensing strategy, the sensitivity primarily depends on how easily a supercoiled DNA gets converted to the relaxed state. Therefore, if a certain supercoiled plasmid requires fewer number of nicking, that plasmid would likely give a more sensitive detection result. Previous studies show that different circular plasmids have varying linking number/degree of supercoiling and thermodynamic stability at supercoiled state^70–73^. At this point, although we do not know exactly what is unique about ΦX174 when compared with pUC19 and pBR322, we suspect that ΦX174 might require fewer number of nicking to convert it into relaxed form. It is also very likely that an even better performing plasmid DNA than ΦX174 may exist. The search for the best ds plasmid reporter is however beyond the scope of this study. Nevertheless, there are a few potential strategies to make this ratiometric sensing strategy more efficient in future. For instance, using DNA topoisomerases (e.g., Topo I, DNA gyrase, Topo III, and Topo IV), we can change the linking number of plasmids to get the best supercoil state that would require the least number of cutting to convert it into relaxed conformation^71^. Also, DNA gyrase enzyme could be utilized to increase the amount of supercoil DNA in the initial sample^74^, which will help set up a high starting ratiometric value for the best contrast.

Next, we compared our results with a typical fluorescent reporting system and found that our ratiometric reporting strategy performed two orders of magnitudes better than the widely used F-Q reporters (**Figure S4**). Besides, this ratiometric sensing strategy is advantageous from a few experimental perspectives while comparing it to our previously demonstrated sizing-based detection: 1) Our previous λ DNA-based reporting system requires a reference lane of unreacted λ DNA to quantify band shift. Still, measurement error could arise because of gel distortion, misalignment, uneven heat distribution and uneven distribution of electric field across gel width and running buffer level^75^. Instead, in our current plasmid-based reporting system, we measure ratiometric band intensity instead of band position, which is independent of gel distortion, misalignment, and running buffer level. 2) In most of the assay methods, including our previous sizing-based signaling mechanism, errors may also arise from the pipetting when adding reporter molecules. Our new strategy is completely immune from this type of human error, because the intensity ratio of supercoiled to relaxed DNA is independent of the absolute amount of reporter molecules added to each reaction. For a thorough comparison, we have included a comprehensive summary table (**Table S2**), juxtaposing our current reporting strategy with established F-Q ssDNA reporters, F-Q dsDNA reporters, and various DNA sizing-based reporters. Notably, we found that the DNA supercoil relaxation technique excels in terms of cost, speed, sensitivity, and ease of signal quantification. Furthermore, DNA supercoil relaxation can easily be evaluated using a more miniaturized microfluidic electrophoresis system^76,77^ instead of standard gel electrophoresis, showing the potential of this method for future POC use like our previous sizing-based methods.

### Validation of ratiometric sensing strategy using real-life targets

Last but not least, we used ΦX174 as the reporter molecule and checked if the supercoil relaxation reporting mechanism could be applied to other DNA targets. First, we checked if the rCRISPR assay is also valid for dsDNA targets. For that, we used a 89-bp long dsDNA as the target (see the sequences in **Table S1**) and ran the rCRISPR assay. The gel image in **Figure S5** confirmed that supercoil relaxation of ΦX174 is equally valid for dsDNA targets. Then, we chose two real-life viral genomes AAV^78^ and HPV 16^47^ and used their genetic markers (see the sequences in **Table S1**) as the model targets to challenge rCRISPR. The gel image in Figure 7 shows that DNA supercoil relaxation mechanism is universally effective for both targets. These results suggest that our simple, ratiometric sensing platform has the potential to detect various genes and pathogens.

**Figure 7.**
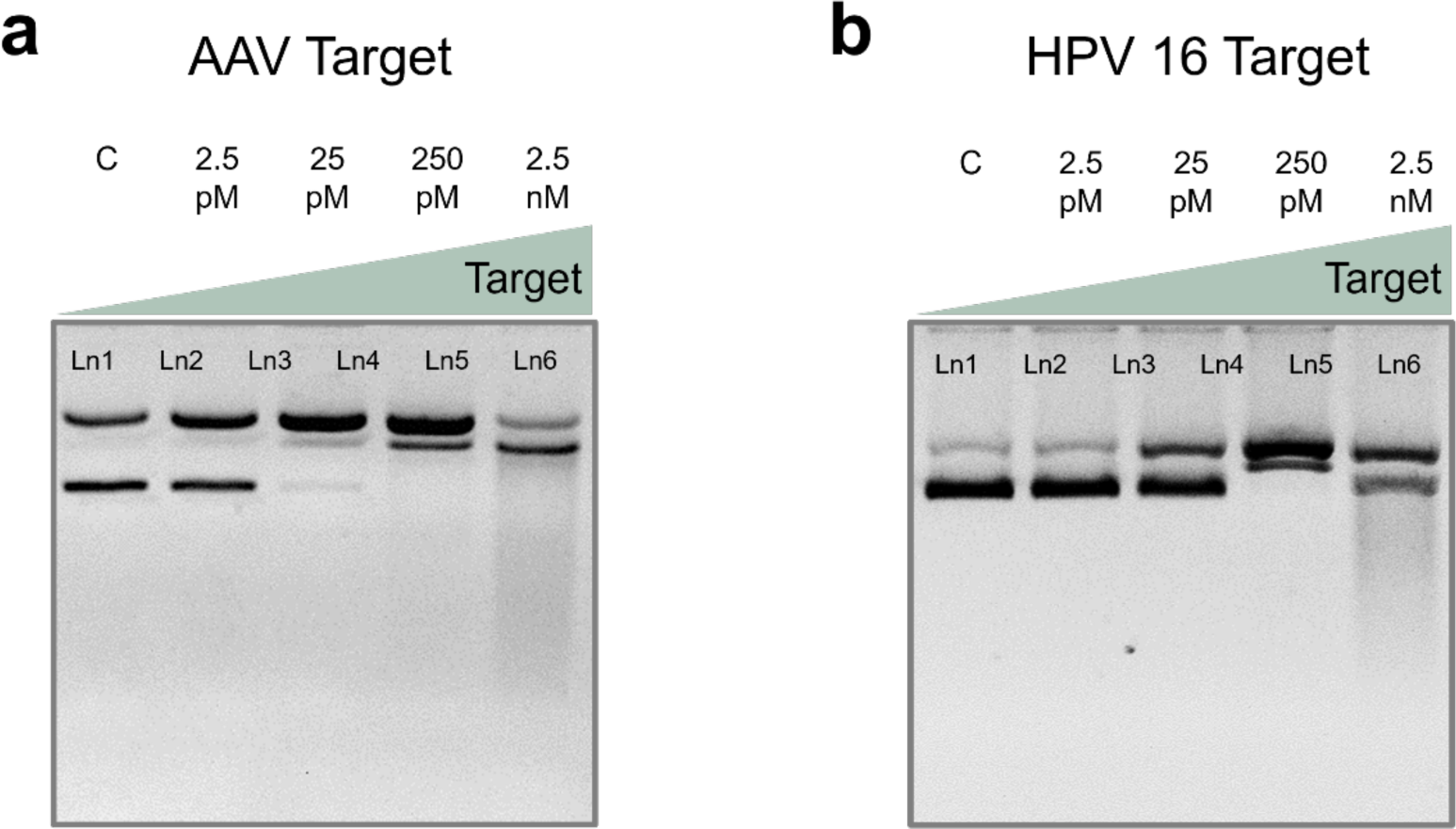
Validation of CRISPR-Cas12a based DNA supercoil relaxation using various targets. Gel electrophoresis (1% agarose gel and 1×TBE buffer) demonstrates the nonspecific supercoil relaxation of ΦX174 DNA. (a) Marker of AAV genome was used as target. (b) Marker of HPV 16 was used as target. In each panel in (a) and (b), Ln1: negative control (no target); Ln2, Ln3, Ln4, Ln5, and Ln6: experimental lanes with target concentrations of 2.5 pM, 25 pM, 250 pM, and 2.5 nM, respectively. Abbreviations: c, negative control; pM, picomolar; nM, nanomolar; Ln, lane; TBE, Tris-borate EDTA. AAV, Adeno-associated virus; HPV, human papillomavirus.

## Conclusions

In summary, we demonstrated rCRISPR, a new strategy for CRISPR-Cas12a-based biosensing by using supercoiled plasmid DNA as ratiometric reporter molecules. We discovered that even though the *trans*-degradation of ds plasmid DNA with activated Cas12a is very slow (>90 min and requires high target concentrations), the *trans*-nicking of supercoil DNA and converting it to relaxed circular conformation is a much faster process (<10 min). This interesting *trans*-nicking property has been leveraged to construct a nonfluorescent, molecular conformation-based CRISPR detection system free of expensive reporter molecules and optical readout equipment. We tested a few dsDNA plasmids, such as pUC19, pBR322, and ΦX174, and found that the DNA supercoil relaxation mechanism is widely prevalent, which makes the new sensing strategy universal. Among these three plasmids, ΦX174 performed the best, which was able to detect ssDNA target with a LOD of ∼250 fM without any preamplification step, two orders better than a typical fluorescent reporting system. Additionally, this ratiometric signaling mechanism obviates all limitations of the previously demonstrated sizing-based signal acquisition technique without compromising sensitivity. As a proof-of-concept, we also applied DNA supercoil relaxation to detect AAV and HPV 16 as model targets. This straightforward yet sensitive strategy is compatible with low-cost and compact readout systems such as microfluidic/mini-gel electrophoresis, making it appealing for POC implementation in the future.

## Methods

### Chemicals and apparatus

LbaCas12a, crRNA, synthetic target, and crRNA were purchased from Integrated DNA Technologies (IDT, Coralville, IA, USA). rCutSmart™ buffer, and NEBuffer™ 2.1, pUC19 DNA (Catalog# FERSD0061), pBR322 DNA (Catalog# FERSD0041 and phiX174 DNA (Catalog# FERSD0031), were purchased from Thermo-fisher. Ultrapure water (18.3 MΩ cm) was produced by the Milli-Q system (Millipore, Inc., USA) and used throughout the experiments. The sequences of oligonucleotides and crRNA molecules are listed in Table S1.

### Nonfluorescent LbaCas12a/gRNA assays

All DNA and RNA oligos were stored in 10 mM Tris-HCl buffer (pH 7.5) at 10 μM concentrations in a -20 °C refrigerator and pre-warmed 37 °C for 30 min before mixing. Reporter molecules (e.g., 1kb dsDNA, pUC19 DNA, pBR322, phiX174) were diluted to 100 ng/µL in 10 mM Tris-HCl buffer (pH 7.5), stored at 4°C and used throughout the experiments. For the nonfluorescent CRISPR assay, analyte solutions with different concentrations of ssDNA were added into the CRISPR reaction reagents, consisting of 40 nM LbaCas12a, 20 nM gRNA, 8-10 ng/ µL reporter molecule and 1X reaction buffer (e.g., 50 mM NaCl, 10 mM Tris-HCl, 10 mM MgCl2, 100 µg/ml BSA, pH=7.9 @ 25°C). The total reaction volume was 80 µL.

### Fluorescent readout

For the fluorescent readout system, 600 nM F-Q reporter molecule (see the sequence in **Table S1**) was used instead of dsDNA plasmid. The CRISPR reaction mixture was incubated in a microplate reader (SpectraMax M2, Molecular Devices) at 37 °C for 1 hr with fluorescence measurements (excitation at 490 nm, and emission at 520 nm) taken every 30 s.

### Nonfluorescent readout using gel electrophoresis

Gel electrophoresis was performed for the nonfluorescent reporting system. 1% agarose gels were prepared using 1× TBE (Tris Borate EDTA) and 1× SYBR™ Safe DNA Gel Stain (Thermo-fisher, catalog# S33102). 10 μL of different reaction products with loading dye (5:1, v/v) were added to each well. Electrophoresis was done at 120 V for 30 min in the same buffer at room temperature. Finally, the agarose gels were scanned and recorded by the E-Gel Imager system (Invitrogen, USA).

### Gel result analysis

Gel results were analyzed using ImageJ software. First, the background was removed from the gel image. The longitudinal intensity of each lane was averaged across the width and plotted. Gel intensity was calculated from the distinct peaks for supercoiled and relaxed circular conformations. Ratiometric intensities (aka. Ratio of supercoiled DNA to relaxed circular DNA) were calculated by dividing supercoiled band intensity by relaxed circular band intensity. Finally, the ratiometric intensities were plotted against the target concentration.

### Statistical analysis

We performed the student’s t-test using repeated experimental data (n=3) to calculate the *p* values to estimate the difference between the experimental results and control groups and determined the LOD. The graph shows statistical insignificance at p>0.05 (ns), and statistical significance at p<0.01 (*), p<0.01 (**), and p<0.001(***).

## Supporting Information

The supporting information is available free of charge at

## Author Contributions

Q.W. and N.M. initiated and conceived the project. N.M. performed all the experiments, collected, and analyzed the data. L.T. and S.D. helped run the gel experiments to produce triplicate data. N.M. and Q.W. wrote the manuscript. All the authors helped revise the manuscript.

## Competing Interests

The authors declare no competing financial interest.

## Acknowledgments

The authors sincerely thank the funding support from the National Science Foundation (Award # 1944167).

## Notes

### Competing Interest Statement

The authors have declared no competing interest.

### Summary of Updates

Typo correction in the main text

## References

1 Smyrlaki, I. et al. Massive and rapid COVID-19 testing is feasible by extraction-free SARS-CoV-2 RT-PCR. Nat. commun. 11, 4812–4812, doi:10.1038/s41467-020-18611-5 (2020).

2 Huang, H. S. et al. Multiplex PCR system for the rapid diagnosis of respiratory virus infection: systematic review and meta-analysis. Clin. Microbiol. Infect. 24, 1055–1063, doi:10.1016/j.cmi.2017.11.018 (2018).

3 Huang, H. S. et al. Multiplex PCR system for the rapid diagnosis of respiratory virus infection: systematic review and meta-analysis. Clinical microbiology and infection : the official publication of the European Society of Clinical Microbiology and Infectious Diseases 24, 1055–1063, doi:10.1016/j.cmi.2017.11.018 (2018).

4 Smyrlaki, I. et al. Massive and rapid COVID-19 testing is feasible by extraction-free SARS-CoV-2 RT-PCR. Nature communications 11, 4812, doi:10.1038/s41467-020-18611-5 (2020).

5 Wu, I. C., Liu, W.-C. & Chang, T.-T. Applications of next-generation sequencing analysis for the detection of hepatocellular carcinoma-associated hepatitis B virus mutations. J. Biomed. Sci. 25, 51–51, doi:10.1186/s12929-018-0442-4 (2018).

6 Niemz, A., Ferguson, T. M. & Boyle, D. S. Point-of-care nucleic acid testing for infectious diseases. Trends Biotechnol. 29, 240–250, doi:10.1016/j.tibtech.2011.01.007 (2011).

7 Ouyang, T., Liu, Z., Han, Z. & Ge, Q. MicroRNA detection specificity: recent advances and future perspective. Anal. Chem. 91, 3179–3186 (2019).

8 Cong, L. et al. Multiplex genome engineering using CRISPR/Cas systems. Science 339, 819–823 (2013).

9 Ledford, H. & Callaway, E. Pioneers of revolutionary CRISPR gene editing win chemistry Nobel. Nature 586, 346–347 (2020).

10 Kim, J. & Kim, J.-S. Bypassing GMO regulations with CRISPR gene editing. Nat. Biotechnol. 34, 1014–1015, doi:10.1038/nbt.3680 (2016).

11 Cyranoski, D. CRISPR gene-editing tested in a person for the first time. Nature 539 (2016).

12 Chen, J. S. et al. CRISPR-Cas12a target binding unleashes indiscriminate single-stranded DNase activity. Science 360, 436–439, doi:10.1126/science.aar6245 (2018).

13 Gootenberg, J. S. et al. Nucleic acid detection with CRISPR-Cas13a/C2c2. Science (New York, N.Y.) 356, 438–442, doi:10.1126/science.aam9321 (2017).

14 Kaminski, M. M., Abudayyeh, O. O., Gootenberg, J. S., Zhang, F. & Collins, J. J. CRISPR-based diagnostics. Nat. Biomed. Eng. 5, 643–656, doi:10.1038/s41551-021-00760-7 (2021).

15 Weng, Z. et al. CRISPR-Cas Biochemistry and CRISPR-Based Molecular Diagnostics. Angew. Chem. Int. Ed. (2023).

16 Mohammad, N., Katkam, S. S. & Wei, Q. Recent Advances in CRISPR-Based Biosensors for Point-of-Care Pathogen Detection. CRISPR J. 5, 500–516 (2022).

17 Mohammad, N., Katkam, S. S. & Wei, Q. A Sensitive and Nonoptical CRISPR Detection Mechanism by Sizing Double-Stranded λ DNA Reporter. Angew. Chem. Int. Ed. 61, e202213920, doi:10.1002/anie.202213920 (2022).

18 Ackerman, C. M. et al. Massively multiplexed nucleic acid detection with Cas13. Nature 582, 277–282, doi:10.1038/s41586-020-2279-8 (2020).

19 Ali, Z. et al. iSCAN: An RT-LAMP-coupled CRISPR-Cas12 module for rapid, sensitive detection of SARS-CoV-2. Virus Res. 288, 198129–198129, doi:10.1016/j.virusres.2020.198129 (2020).

20 Arizti-Sanz, J. et al. Equipment-free detection of SARS-CoV-2 and Variants of Concern using Cas13. medRxiv, 2021.2011.2001.21265764, doi:10.1101/2021.11.01.21265764 (2021).

21 Arizti-Sanz, J. et al. Streamlined inactivation, amplification, and Cas13-based detection of SARS-CoV-2. Nat. Commun. 11, 5921, doi:10.1038/s41467-020-19097-x (2020).

22 Bai, J. et al. Cas12a-Based On-Site and Rapid Nucleic Acid Detection of African Swine Fever. Front. Microbiol. 10, 2830–2830, doi:10.3389/fmicb.2019.02830 (2019).

23 Barnes, K. G. et al. Deployable CRISPR-Cas13a diagnostic tools to detect and report Ebola and Lassa virus cases in real-time. Nat. Commun. 11, 4131–4131, doi:10.1038/s41467-020-17994-9 (2020).

24 Broughton, J. P. et al. CRISPR–Cas12-based detection of SARS-CoV-2. Nat. Biotechnol. 38, 870–874, doi:10.1038/s41587-020-0513-4 (2020).

25 Chaijarasphong, T., Thammachai, T., Itsathitphaisarn, O., Sritunyalucksana, K. & Suebsing, R. Potential application of CRISPR-Cas12a fluorescence assay coupled with rapid nucleic acid amplification for detection of white spot syndrome virus in shrimp. Aquaculture 512, 734340, doi:10.1016/j.aquaculture.2019.734340 (2019).

26 Chang, Y. et al. Visual detection of porcine reproductive and respiratory syndrome virus using CRISPR-Cas13a. Transbound. Emerg. Dis. 67, 564–571, doi:10.1111/tbed.13368 (2020).

27 Chen, M. et al. CRISPR/Cas12a-Based Ultrasensitive and Rapid Detection of JAK2 V617F Somatic Mutation in Myeloproliferative Neoplasms. Biosensors 11, 247 (2021).

28 Chen, Y. et al. Contamination-free visual detection of SARS-CoV-2 with CRISPR/Cas12a: A promising method in the point-of-care detection. Biosens. Bioelectron. 169, 112642–112642, doi:10.1016/j.bios.2020.112642 (2020).

29 Dai, Y. et al. Exploring the Trans-Cleavage Activity of CRISPR-Cas12a (cpf1) for the Development of a Universal Electrochemical Biosensor. Angew. Chem. Int. Ed. 58, 17399–17405, doi:10.1002/anie.201910772 (2019).

30 de Puig, H. et al. Minimally instrumented SHERLOCK (miSHERLOCK) for CRISPR-based point-of-care diagnosis of SARS-CoV-2 and emerging variants. Sci. Adv. 7, eabh2944 (2021).

31 Ding, X. et al. Ultrasensitive and visual detection of SARS-CoV-2 using all-in-one dual CRISPR-Cas12a assay. Nat. Commun. 11, 4711–4711, doi:10.1038/s41467-020-18575-6 (2020).

32 Ding, X., Yin, K., Li, Z., Sfeir, M. M. & Liu, C. Sensitive quantitative detection of SARS-CoV-2 in clinical samples using digital warm-start CRISPR assay. Biosens. Bioelectron. 184, 113218–113218, doi:10.1016/j.bios.2021.113218 (2021).

33 Fozouni, P. et al. Amplification-free detection of SARS-CoV-2 with CRISPR-Cas13a and mobile phone microscopy. Cell 184, 323–333.e329, doi:10.1016/j.cell.2020.12.001 (2021).

34 Gootenberg, J. S. et al. Multiplexed and portable nucleic acid detection platform with Cas13, Cas12a, and Csm6. Science (New York, N.Y.) 360, 439–444, doi:10.1126/science.aaq0179 (2018).

35 Guo, L. et al. SARS-CoV-2 detection with CRISPR diagnostics. Cell Discov. 6, 34–34, doi:10.1038/s41421-020-0174-y (2020).

36 Jiao, J. et al. Field detection of multiple RNA viruses/viroids in apple using a CRISPR/Cas12a-based visual assay. Plant Biotechnol. J. 19, 394–405, doi:10.1111/pbi.13474 (2021).

37 Joung, J. et al. Detection of SARS-CoV-2 with SHERLOCK One-Pot Testing. N. Engl. J. Med. 383, 1492–1494, doi:10.1056/NEJMc2026172 (2020).

38 Li, S.-Y. et al. CRISPR-Cas12a-assisted nucleic acid detection. Cell Discov. 4, 20–20, doi:10.1038/s41421-018-0028-z (2018).

39 Mustafa, M. I. & Makhawi, A. M. SHERLOCK and DETECTR: CRISPR-Cas Systems as Potential Rapid Diagnostic Tools for Emerging Infectious Diseases. J. Clin. Microbiol. 59, e00745–00720, doi:10.1128/JCM.00745-20 (2021).

40 Myhrvold, C. et al. Field-deployable viral diagnostics using CRISPR-Cas13. Science (New York, N.Y.) 360, 444–448, doi:10.1126/science.aas8836 (2018).

41 Nguyen, L. T., Smith, B. M. & Jain, P. K. Enhancement of trans-cleavage activity of Cas12a with engineered crRNA enables amplified nucleic acid detection. Nat. Commun. 11, 4906–4906, doi:10.1038/s41467-020-18615-1 (2020).

42 Palaz, F., Kalkan, A. K., Tozluyurt, A. & Ozsoz, M. CRISPR-based tools: Alternative methods for the diagnosis of COVID-19. Clin. Biochem. 89, 1–13, doi:10.1016/j.clinbiochem.2020.12.011 (2021).

43 Park, J. S. et al. Digital CRISPR/Cas-Assisted Assay for Rapid and Sensitive Detection of SARS-CoV-2. Adv. Sci. (Weinheim, Baden-Wurttemberg, Germany) 8, 2003564–2003564, doi:10.1002/advs.202003564 (2021).

44 Patchsung, M. et al. Clinical validation of a Cas13-based assay for the detection of SARS-CoV-2 RNA. Nat. Biomed. Eng. 4, 1140–1149, doi:10.1038/s41551-020-00603-x (2020).

45 Rahimi, H. et al. CRISPR Systems for COVID-19 Diagnosis. ACS Sens. 6, 1430–1445, doi:10.1021/acssensors.0c02312 (2021).

46 Shi, K. et al. A CRISPR-Cas autocatalysis-driven feedback amplification network for supersensitive DNA diagnostics. Sci. Adv. 7, eabc7802, doi:10.1126/sciadv.abc7802 (2021).

47 Yu, T. et al. Coupling smartphone and CRISPR–Cas12a for digital and multiplexed nucleic acid detection. AIChE J. n/a, e17365, doi:10.1002/aic.17365 (2021).

48 Ning, B. et al. A smartphone-read ultrasensitive and quantitative saliva test for COVID-19. Sci. Adv.7, eabe3703 (2021).

49 Wang, X. et al. CRISPR/Cas12a technology combined with immunochromatographic strips for portable detection of African swine fever virus. Commun. Biol. 3, 62–62, doi:10.1038/s42003-020-0796-5 (2020).

50 Li, Y., Mansour, H., Wang, T., Poojari, S. & Li, F. Naked-Eye Detection of Grapevine Red-Blotch Viral Infection Using a Plasmonic CRISPR Cas12a Assay. Anal. Chem. 91, 11510–11513, doi:10.1021/acs.analchem.9b03545 (2019).

51 Yan, H. et al. A one-pot isothermal Cas12-based assay for the sensitive detection of microRNAs. Nat Biomed Eng, 1–19 (2023).

52 Rananaware, S. R. et al. Programmable RNA detection with CRISPR-Cas12a. bioRxiv, 2023–2001 (2023).

53 Pang, Y. et al. CRISPR-cas12a mediated SERS lateral flow assay for amplification-free detection of double-stranded DNA and single-base mutation. Chem. Eng. J. 429, 132109, doi:10.1016/j.cej.2021.132109 (2022).

54 Hu, M. et al. Single-Step, Salt-Aging-Free, and Thiol-Free Freezing Construction of AuNP-Based Bioprobes for Advancing CRISPR-Based Diagnostics. JACS. 142, 7506–7513, doi:10.1021/jacs.0c00217 (2020).

55 Yuan, C. et al. Universal and Naked-Eye Gene Detection Platform Based on the Clustered Regularly Interspaced Short Palindromic Repeats/Cas12a/13a System. Anal. Chem. 92, 4029–4037, doi:10.1021/acs.analchem.9b05597 (2020).

56 Qin, P. et al. Rapid and Fully Microfluidic Ebola Virus Detection with CRISPR-Cas13a. ACS Sens. 4, 1048–1054, doi:10.1021/acssensors.9b00239 (2019).

57 Xu, W., Jin, T., Dai, Y. & Liu, C. C. Surpassing the detection limit and accuracy of the electrochemical DNA sensor through the application of CRISPR Cas systems. Biosens. Bioelectron. 155, 112100, doi:10.1016/j.bios.2020.112100 (2020).

58 Dai, Y. et al. Exploring the Trans-Cleavage Activity of CRISPR-Cas12a (cpf1) for the Development of a Universal Electrochemical Biosensor. Angewandte Chemie (International ed. in English) 58, 17399–17405, doi:10.1002/anie.201910772 (2019).

59 Birren, B. W., Lai, E., Hood, L. & Simon, M. I. Pulsed field gel electrophoresis techniques for separating 1-to 50-kilobase DNA fragments. Anal. Biochem. 177, 282–286 (1989).

60 Herschleb, J., Ananiev, G. & Schwartz, D. C. Pulsed-field gel electrophoresis. Nat. Protoc. 2, 677–684 (2007).

61 Mohammad, N., Talton, L., Hetzler, Z., Gongireddy, M. & Wei, Q. Unidirectional trans-cleaving behavior of CRISPR-Cas12a unlocks for an ultrasensitive assay using hybrid DNA reporters containing a 3’ toehold. Nucleic Acids Res. 51, 9894–9904 (2023).

62 Yanisch-Perron, C., Vieira, J. & Messing, J. Improved M13 phage cloning vectors and host strains: nucleotide sequences of the M13mpl8 and pUC19 vectors. Gene 33, 103–119 (1985).

63 Betlach, M. C., Heynecker, H. L. & Boyer, H. W. pBR322 DNA. Gene 2, 95–113 (1977).

64 Sanger, F. et al. Nucleotide sequence of bacteriophage ΦX174 DNA. nature 265, 687–695 (1977).

65 Murugan, K., Seetharam, A. S., Severin, A. J. & Sashital, D. G. CRISPR-Cas12a has widespread off-target and dsDNA-nicking effects. JBC. 295, 5538–5553 (2020).

66 Matange, K., Tuck, J. M. & Keung, A. J. DNA stability: a central design consideration for DNA data storage systems. Nat. Commun. 12, 1358, doi:10.1038/s41467-021-21587-5 (2021).

67 Wang, X., Lim, H. J. & Son, A. Characterization of denaturation and renaturation of DNA for DNA hybridization. Environ. Toxicol. 29 (2014).

68 Nguyen, L. T. et al. Engineering highly thermostable Cas12b via de novo structural analyses for one-pot detection of nucleic acids. Cell Rep. 4 (2023).

69 Tan, Z. J. & Chen, S. J. Nucleic acid helix stability: effects of salt concentration, cation valence and size, and chain length. Biophys. J. 90, 1175–1190, doi:10.1529/biophysj.105.070904 (2006).

70 Xu, Y. C. & Qian, L. Determination of linking number of pBR322 DNA. Sci. Sin. B 26, 602–613 (1983).

71 Higgins, N. P. & Vologodskii, A. V. Topological behavior of plasmid DNA. Microbiol. Spectr. 3, 3–2 (2015).

72 Serban, D., Benevides, J. M. & Thomas, G. J., Jr. DNA secondary structure and Raman markers of supercoiling in Escherichia coli plasmid pUC19. Biochem. 41, 847–853, doi:10.1021/bi011004z (2002).

73 Vologodskii, A. V., Levene, S. D., Klenin, K. V., Frank-Kamenetskii, M. & Cozzarelli, N. R. Conformational and thermodynamic properties of supercoiled DNA. J. Mol. Biol. 227, 1224–1243, doi:10.1016/0022-2836(92)90533-P (1992).

74 Cozzarelli, N. R. DNA gyrase and the supercoiling of DNA. Science 207, 953–960 (1980).

75 Scientific, T. F. Nucleic Acid Electrophoresis Education.

76 Gutzweiler, L. et al. Open microfluidic gel electrophoresis: Rapid and low cost separation and analysis of DNA at the nanoliter scale. Electrophor. 38, 1764–1770, doi:10.1002/elps.201700001 (2017).

77 Duong, T. T. et al. Size-dependent free solution DNA electrophoresis in structured microfluidic systems. Microelectron. Eng. 67-68, 905–912, doi:10.1016/S0167-9317(03)00153-9 (2003).

78 Hetzler, Z. et al. Rapid Adeno-Associated Virus Genome Quantification with Amplification-Free CRISPR-Cas12a. bioRxiv, 2023–2011 (2023).

