## Supporting information for "A Ratiometric Nonfluorescent CRISPR Assay Utilizing Cas12a-Induced Plasmid Supercoil Relaxation"

**Optimization: Effect of reaction time on trans-nicking of pUC19**

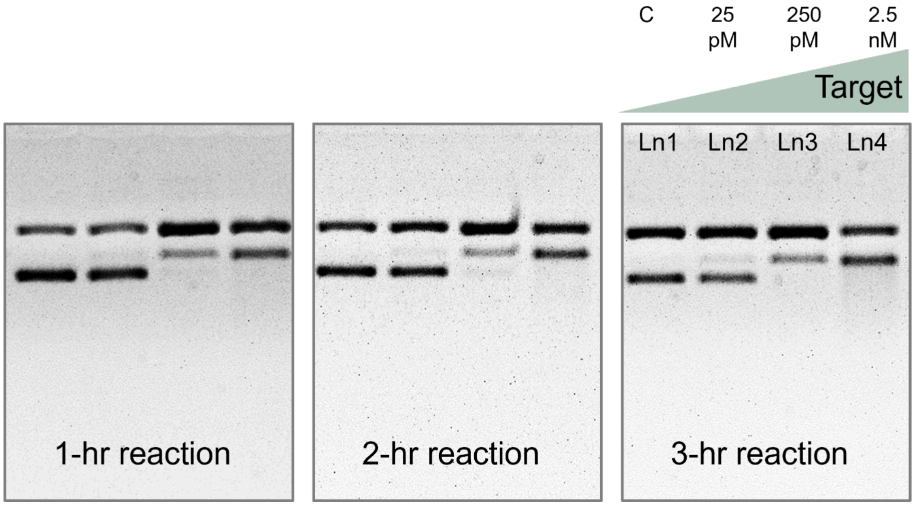


**Figure S1**. Gel electrophoresis (1% agarose gel and 1×TBE buffer) results demonstrating the effect of reaction time on trans-nicking of pUC19 reporters. Abbreviations: c, negative control; pM, picomolar; nM, nanomolar; TBE, Tris-borate EDTA.

**Intensity diagram for optimization of pretreatment and reaction temperature**

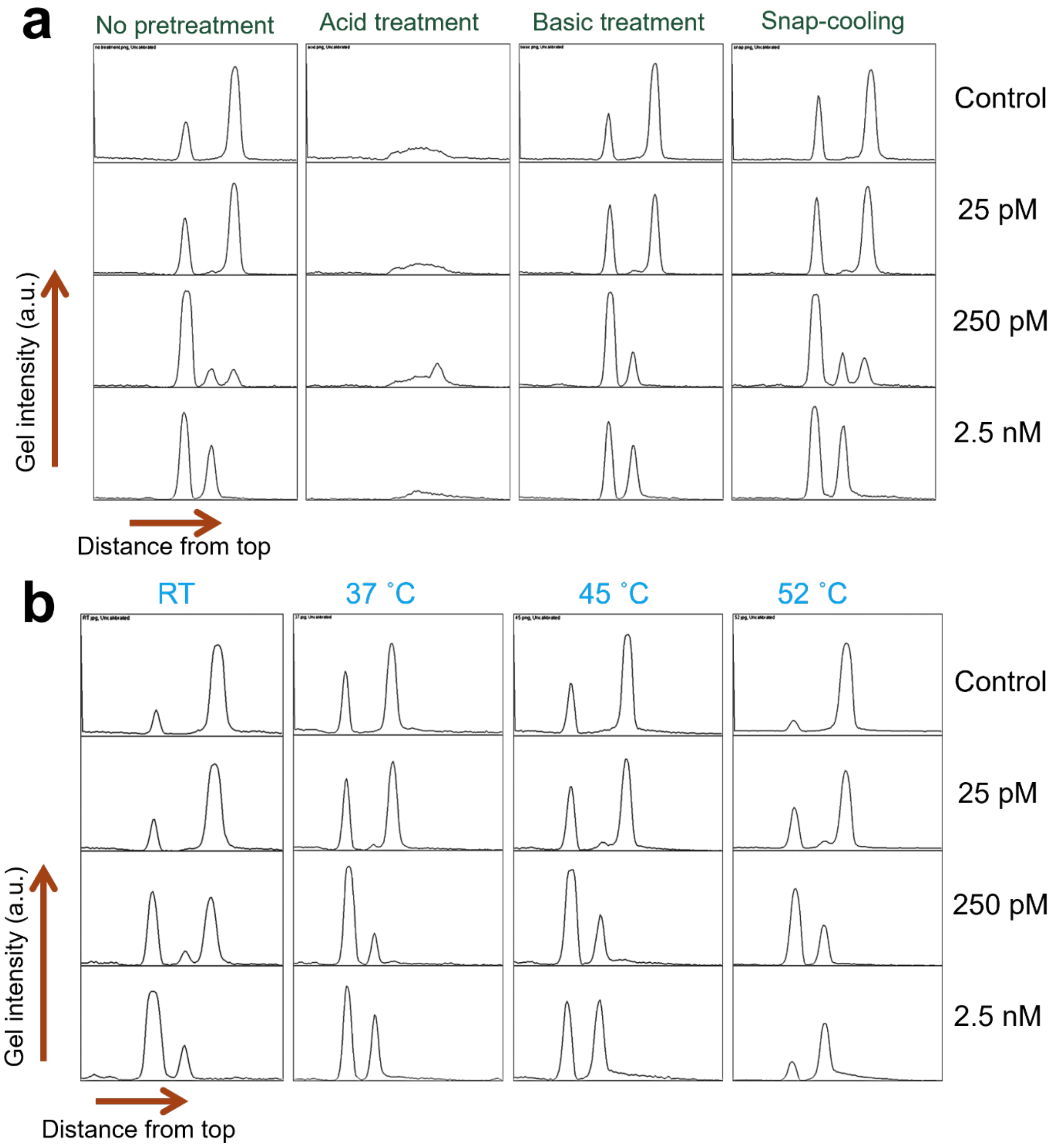


**Figure S2**. Intensity diagrams of gel results to show the effect of (a) pretreatment of reporter molecule, and (b) reaction temperature. Abbreviations: c, negative control; pM, picomolar; nM, nanomolar; RT, room temperature; a.u., arbitrary unit.

### Intensity diagram for optimization of Cas12a concentration, buffer type and salt concentration

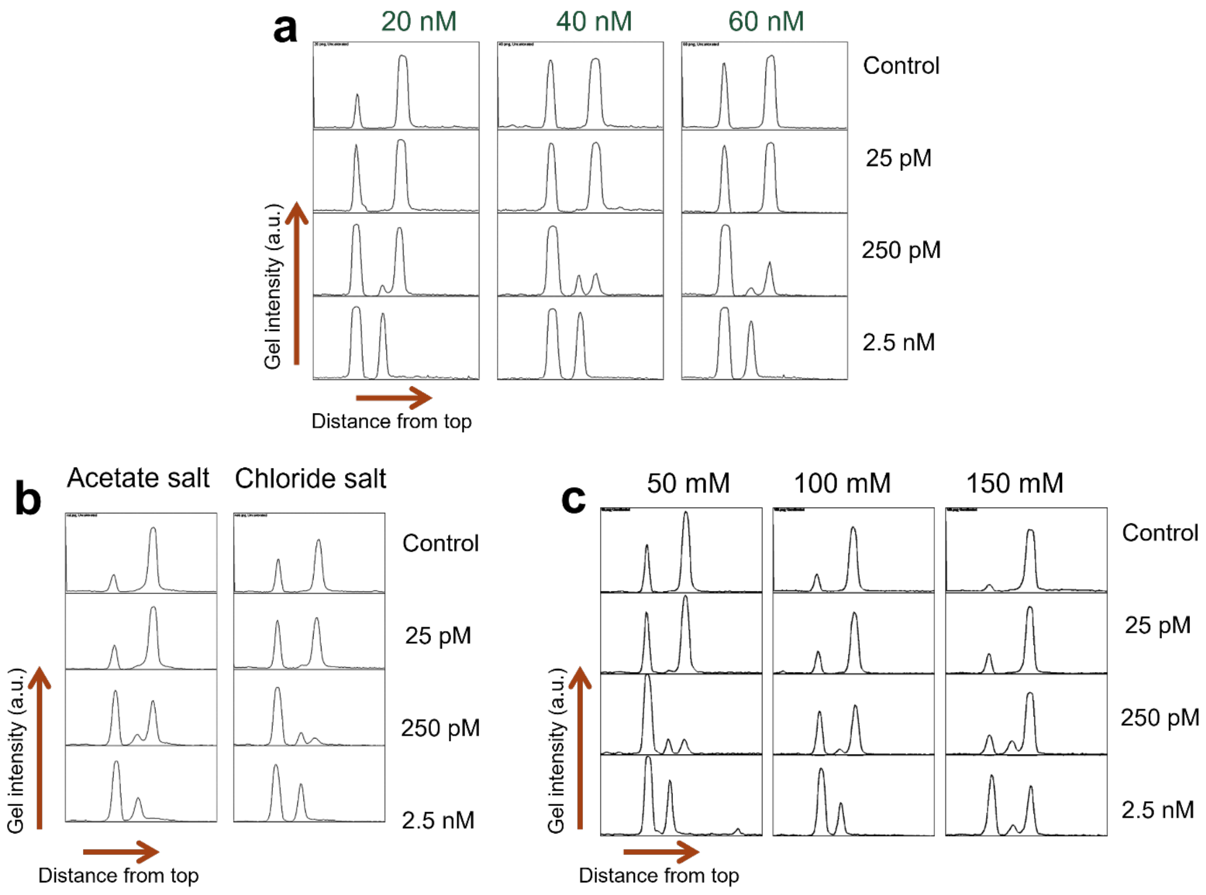


**Figure S3**. Intensity diagrams of gel results to show the effect of (a) Cas12a concentration, (b) buffer type, and (c) salt concentration. Abbreviations: c, negative control; pM, picomolar; nM, nanomolar; mM, millimolar; a.u., arbitrary unit.

### CRISPR-Cas12a based detection using conventional fluorescent reporters

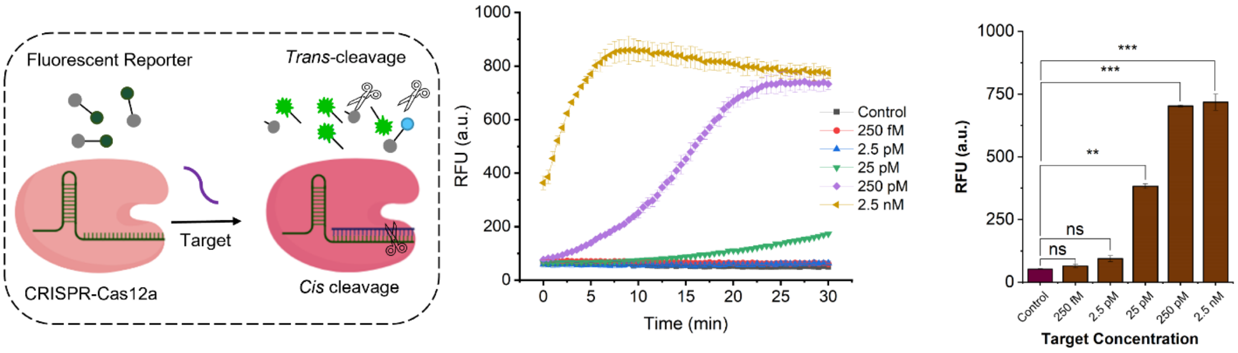


**Figure S4**. CRISPR-LbCas12a based detection using conventional fluorescent reporter molecules. (a) schematic diagram of CRISPR-LbCas12a assay. (b) RFU signal acquired for 30 min reaction with different target concentrations. (c) RFU reading taken after 1 hr of reaction against different target concentrations. The graph shows statistical insignificance at p>0.05 (ns); and statistical significance at p<0.01 (**), and p<0.001(***). Abbreviations: RFU, relative fluorescence unit, ns, not significant.

**Verifying ratiometric sensing strategy using dsDNA target**

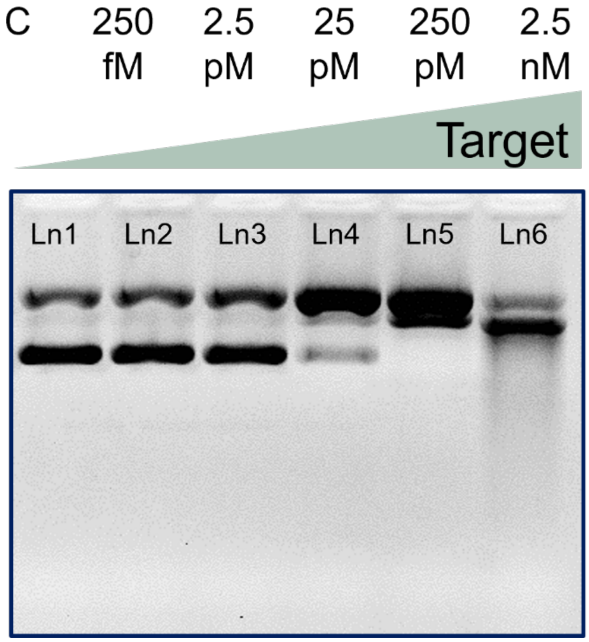


**Figure S5**. Validation of rCRISPR for detecting dsDNA targets. Gel electrophoresis (1% agarose gel and 1×TBE buffer) results demonstrating the nonspecific supercoil relaxation of ΦX174 DNA. Ln1: negative control (no target); Ln2, Ln3, Ln4, Ln5 and Ln6: experimental lanes with target concentrations of 250 fM, 2.5 pM, 25 pM, 250 pM, and 2.5 nM, respectively. Abbreviations: c, negative control; fM, femtomolar; pM, picomolar; nM, nanomolar; Ln, lane; TBE, Tris-borate EDTA.

**Table S1. Oligos used for the study.**

| Item | Sequence (5'-3') |
|---|---|
| gRNA | UAAUUUCUACUAAGUGUAGAUCGUCGCCGUCCAGCUCGACC |
| Target ssDNA | GGT CGA GCT GGA CGG CGA CG |
| Target dsDNA | Sense strand: GGT CGA GCT GGA CGG CGA CGT AAA TCG ACG ACG CTG ACG GTA GCG AAT CGA TCG TAC GCT AGT CCG TAA TGT GAG TTG GCT GAT GGT TA<br>Antisense strand: TAA CCA TCA GCC AAC TCA CAT TAC GGA CTA GCG TAC GAT CGA TTC GCT ACC GTC AGC GTC GTC GAT TTA CGT CGC CGT CCA GCT CGA CC |
| gRNA_AAV | UA AUU UCU ACU AAG UGU AGA UCU CCA UCA CUA GGG GUUCCU |
| Target AAV | AGG AAC CCC TAG TGA TGG AG |
| gRNA_HPV16 | UAA UUU CUA CUC UUG UAG AUU GAA GUA GAU AUG GCAGCAC |
| Target HPV16 | GTG CTG CCA TAT CTA CTT CA |
| F-Q reporter | /6-FAM/ AAAAA /Dabcyl/ |

**Table S2. Performance comparison of various reporters**

| Reporting types | | LOD (pM) | Relative reaction rate | Easy of signal quantification | Relative Cost | Ref. |
|---|---|---|---|---|---|---|
| F-Q ssDNA reporter | 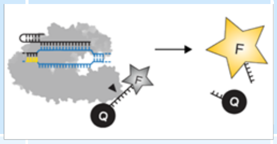  | ~10 | High | Easy | High | [1] |
| F-Q dsDNA reporter (<30bp) | 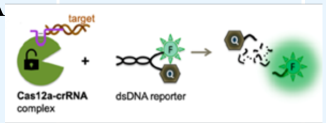  | ~10 | Low | Easy | High | [2] |
| Long dsDNA reporter (sizing-based detection) | 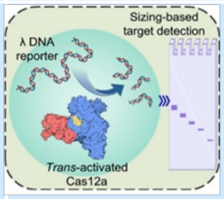  | ~0.25 | High | Hard | Low | Mohammad et al. [3] |
| Hybrid reporter | 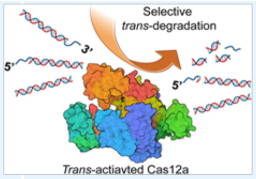  | ~0.25 | Low | Easy | Low | Mohammad et al. [4] |
| DNA supercoil relaxation | 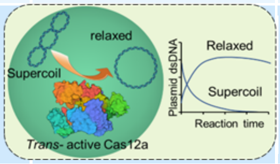  | ~0.25 | High | Easy | Low | This research |

References:


1 Chen, J. S. *et al.* CRISPR-Cas12a target binding unleashes indiscriminate single-stranded DNase activity. *Science* **360**, 436-439, doi:10.1126/science.aar6245 (2018).

2 Smith, C. W. *et al.* Probing CRISPR-Cas12a Nuclease Activity Using Double-Stranded DNA-Templated Fluorescent Substrates. *Biochem.* **59**, 1474-1481, doi:10.1021/acs.biochem.0c00140 (2020).

3 Mohammad, N., Katkam, S. S. & Wei, Q. A Sensitive and Nonoptical CRISPR Detection Mechanism by Sizing Double-Stranded λ DNA Reporter. *Angew. Chem. Int. Ed.* **61**, e202213920, doi:https://doi.org/10.1002/anie.202213920 (2022).

4 Mohammad, N., Talton, L., Hetzler, Z., Gongireddy, M. & Wei, Q. Unidirectional trans-cleaving behavior of CRISPR-Cas12a unlocks for an ultrasensitive assay using hybrid DNA reporters containing a 3′ toehold. *Nucleic Acids Res.* **51**, 9894-9904 (2023).
